# Travelling waves due to negative plant-soil feedbacks in a model including tree life-stages

**DOI:** 10.1101/2023.06.09.544359

**Authors:** Annalisa Iuorio, Mara Baudena, Maarten B. Eppinga, Francesco Giannino, Max Rietkerk, Frits Veerman

**Affiliations:** University of Vienna, Faculty of Mathematics, Oskar-Morgenstern-Platz 1, Vienna, 1090, Austria; Utrecht University, Copernicus Institute of Sustainable Development, Environmental Sciences Group, Utrecht, 3508 TC, The Netherlands; National Research Council of Italy, Institute of Atmospheric Sciences and Climate (CNR-ISAC), Corso Fiume 4, Torino, 10133, Italy; University of Zurich, Department of Geography, Winterthurerstrasse 190, Zürich, 8057, Switzerland; University of Naples Federico II, Department of Agricultural Sciences, via Università 100, Portici, 80055, Italy; Leiden University, Mathematical Institute, Niels Bohrweg 1, Leiden, 2300 RA, The Netherlands

**Keywords:** reaction-diffusion-ODE, Janzen-Connell hypothesis, autotoxicity, travelling waves, linear spreading speed, negative feedback

## Abstract

The emergence and maintenance of tree species diversity in tropical forests is commonly attributed to the Janzen-Connell (JC) hypothesis, which states that growth of seedlings is suppressed in the proximity of conspecific adult trees. As a result, a JC distribution due to a density-dependent negative feedback emerges in the form of a (transient) pattern where conspecific seedling density is highest at intermediate distances away from parent trees. Several studies suggest that the required density-dependent feedbacks behind this pattern could result from interactions between trees and soil-borne pathogens. However, negative plant-soil feedback may involve additional mechanisms, including the accumulation of autotoxic compounds generated through tree litter decomposition. An essential task therefore consists in constructing mathematical models incorporating both effects showing the ability to support the emergence of JC distributions.

In this work, we develop and analyse a novel reaction-diffusion-ODE model, describing the interactions within tropical tree species across different life stages (seeds, seedlings, and adults) as driven by negative plant-soil feedback. In particular, we show that under strong negative plant-soil feedback travelling wave solutions exist, creating transient distributions of adult trees and seedlings that are in agreement with the Janzen-Connell hypothesis. Moreover, we show that these travelling wave solutions are pulled fronts and a robust feature as they occur over a broad parameter range. Finally, we calculate their linear spreading speed and show its (in)dependence on relevant nondimensional parameters.

**2020 MSC:** 35C07, 34C60, 34D05, 35K57, 37C25, 65M06, 92D40.

## 1. Introduction

A widely observed phenomenon in forest tree communities is that conspecific seedling density is highest at intermediate distances from the parent tree, referred to as the Janzen-Connell (JC) distribution. The emergence of JC distributions provide an explanation for the creation and maintenance of high species diversity in forest tree communities [7, 14]. This (transient) pattern is particularly important in terms of biodiversity: the space between the parent tree and its seedlings is a favourable area for other species to colonise and grow, enhancing coexistence (see e.g. [20, 27]). From an ecological viewpoint, an increasing number of ecological studies is supporting the idea that the emergence of this pattern (particularly prominent in tropical ecosystems) is strongly linked to negative plant-soil feedbacks [1, 19, 32]. Among the main mechanisms responsible for such feedbacks, the accumulation of species-specific soil pathogens is indicated as prominent [1, 21]. Consequently, several models have been introduced in the last few decades to theoretically investigate this mechanism (see e.g. [25, 26, 31, 35] and references therein). In recent years, additional mechanisms generating negative plant-soil feedback have been identified, including the accumulation of conspecific extracellular DNA fragments leading to an autotoxic soil environment [4, 30]. Such negative feedback induced by autotoxicity could potentially explain species coexistence in diverse communities [2, 28, 29] as well as plants spatial organisation by means of as clonal rings [3, 6], fairy rings [15, 34], and more generally vegetation patterns [22, 23].

In this work, we construct a new model based on reaction-diffusion-ODEs in order to describe the emergence of JC distributions including both growth inhibition (induced by extracellular self-DNA) and increased mortality (mainly linked to the accumulation of soil-borne pathogens). Reaction-diffusion-ODE systems are used to model a wide variety of phenomena in biology; however, only few analytical results concerning their behaviour – which often strongly differs from classical reaction-diffusion models – are available, see e.g. [11, 13, 16, 24, 37].

As both growth inhibition and increased mortality mechanisms act on different tree life-stages, we consider a stage-structured framework. Our aim consists in introducing a theoretical tool which may help assessing the relative contribution of both mechanisms to emergent spatial distributions of adult trees and their seedlings. As JC distributions are experimentally observed as transient patterns, we analytically investigate the existence of travelling wave solutions which exhibit the typical JC feature of seedlings’ biomass being at a maximum at suitable distances from the parent tree. Travelling wave solutions are widely found in mathematical models inspired by several biological applications, including e.g. species competition [5], tumour growth [9], and bacterial chemotaxis [10]. In particular, we show the existence of such solutions and derive corresponding relevant properties. Moreover, we hypothesize that the constructed travelling wave solutions correspond to pulled fronts, whose speed then coincides with the linear speed determined by a linear analysis near the trivial steady state. We then analytically derive the linear speed and confirm our prediction by comparing the analytical value with the one obtained by numerical simulations of our model for a set of fixed parameter values and investigating their dependence with respect to two relevant parameters.

The impact of the work presented here is twofold: from the ecological viewpoint, our work provides a valuable theoretical tool to further address relevant issues related to JC distributions (e.g. understanding how the dispersal ability of tree species moderate the spatial patterns of adult and seedlings and to what extent are plant strategies along the growth-defence trade-off reflected in the spatial patterns of adult and seedlings). From the mathematical viewpoint, on the other hand, the analytical strategy used here to investigate travelling waves in a system of 4 reaction-diffusion-ODEs improves our understanding of such complex systems and offers a framework potentially useful to investigate problems exhibiting a similar structure.

The paper is structured as follows: in Section 2 we introduce the model both in its dimensional and nondimensional form, on which we focus for our subsequent analysis. The spatially homogeneous steady states associated to this model are derived in Section 3. In Section 4 the linear stability of these steady states with respect to both homogeneous and heterogeneous perturbations is carried out, revealing the absence of Turing patterns for the parameter ranges defined based on experimental findings (as expected). The existence and the main properties of travelling wave solutions (in particular right-moving fronts) are then investigated in Section 5: numerical simulations suggesting the existence of pulled fronts are corroborated analytically by deriving the linear wave speed and comparing it with the numerical measured speed. We conclude our work with a discussion of the results obtained and an outlook indicating further research perspectives in Section 6.

## 2. The model

In our framework, negative plant-soil feedback (NF) manifests itself both during the seed-to-seedling transition (in terms of *growth inhibition*) and at the seedlings life-stage (in terms of *increased mortality*). The first effect can be often attributed to the presence of extracellular self-DNA (also known as autotoxicity), whereas the second effect is mainly linked to soil-borne pathogens. As these factors act at different stages of a tree lifespan, vegetation is considered in terms of biomass and is divided into three compartments corresponding to three different lifestages, namely seeds *Ŝ* (kg*/*m^2^), seedlings 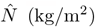 and adults *Â* (kg*/*m^2^). Moreover, the general inhibitor variable *Î* (kg*/*m^2^) represents the density of inhibitor responsible both for growth inhibition and increased mortality effects. The interaction of such variables at any spatial point 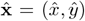 and any time 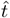 is based on the following assumptions: the increase of seed density is influenced by adult tree production via the per capita seed production rate *ĝ*_*S*_ and seed dispersal 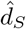, whereas their natural decay rate (including predation) is represented by 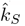. Seeds then germinate and the seedlings might establish or not, depending also on the inhibitor due to the effect of autotoxicity via the function 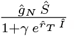. Seedlings have a natural turnover rate 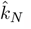, enhanced by pathogens via the term 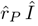. The seedlings which survive then grow into the next life stage according to the function 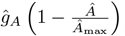. Adults’ density grows logistically because of seedlings transitioning to the adult stage at rate *ĝ*_*A*_, intrinsic growth rate *ĉ*_*A*_, and constant per capita mortality rate 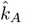. The inhibitor density grows due to adult decomposition byproducts at a rate *ĉ*_*T*_, decays naturally at a rate 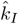, and diffuses in the soil at a rate determined by the coefficient the coefficient 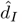. These ecological processes are described by the following reaction-diffusion-ODE system:

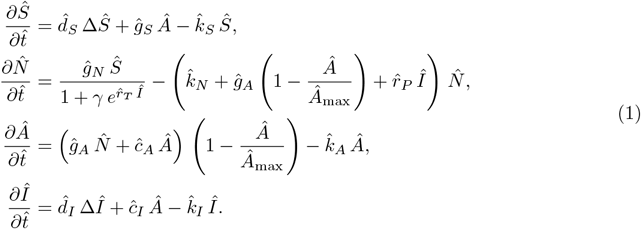

Values and meaning of the non-negative model parameters in (1) are provided in Table 1. Based on an ecological investigation, they have been calibrated in some cases and parametrised in all the others [12].

**Table 1:** Description, values, and units for model parameters in System (1), obtained through parametrisation and calibration.

| Parameter | Description | Value | Units |
| --- | --- | --- | --- |
| $\hat{g}_S$ | Growth rate of $\hat{S}$ | $6.67 \cdot 10^{-8} - 0.033$ | $y^{-1}$ |
| $\hat{k}_S$ | $\hat{S}$ turnover rate | $0.33 - 0.5$ | $y^{-1}$ |
| $\hat{g}_N$ | Transition rate from $\hat{S}$ to $\hat{N}$ | $0.25 - 25$ | $y^{-1}$ |
| $\gamma$ | Establishment sensitivity to toxicity parameter | $10^{-5}$ | - |
| $\hat{r}_T$ | Establishment sensitivity to toxicity parameter | $0 - 68$ | $m^2 kg^{-1}$ |
| $\hat{k}_N$ | Death rate of $\hat{N}$ | $0.02 - 0.74$ | $y^{-1}$ |
| $\hat{r}_P$ | Extra mortality of $\hat{N}$ caused by $\hat{I}$ | $0 - 2$ | $m^2 kg^{-1} y^{-1}$ |
| $\hat{g}_A$ | Transition rate from $\hat{N}$ to $\hat{A}$ | $0.02 - 1$ | $y^{-1}$ |
| $\hat{c}_A$ | Growth rate in $\hat{A}$'s biomass density | $0.25$ | $y^{-1}$ |
| $\hat{A}_{\max}$ | Maximum capacity for $\hat{A}$ | $30$ | $m^{-2} kg$ |
| $\hat{k}_A$ | Mortality rate of $\hat{A}$ | $0.01$ | $y^{-1}$ |
| $\hat{c}_I$ | Growth rate of $\hat{I}$ due to $\hat{A}$ | $1$ | $y^{-1}$ |
| $\hat{k}_I$ | Inhibitor decay rate | $0.7$ | $y^{-1}$ |
| $\hat{d}_S$ | Diffusion coefficient for $\hat{S}$ | $3 - 4$ | $m^2 y^{-1}$ |
| $\hat{d}_I$ | Diffusion coefficient for $\hat{I}$ | $0 - 10$ | $m^2 y^{-1}$ |

In this framework, links to the Janzen-Connell hypothesis can be found in transient patterns where a ring of seedlings emerges around the adult tree (whose density is concentrated in the centre of the ring). Mathematically, this consists in travelling wave solutions, whose construction we analyse in this work. From here on, we refer to this phenomenon as the Janzen-Connell distribution.

In order to reduce the total number of parameters and to facilitate the analytical investigation of our model, we introduce a non-dimensional version of System (1). Existence and stability (both under homogeneous and heterogeneous perturbations) of the corresponding steady states are then investigated in Section 3 and 4, respectively.

In order to facilitate the investigation of the existence and stability properties of our model, we introduce the following nondimensional variables:

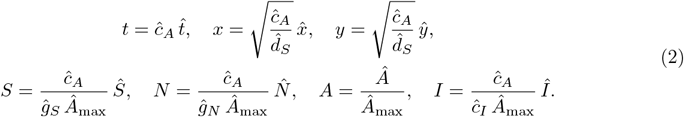

We choose *ĉ*_*A*_, the growth rate of the adult biomass density, as the characteristic time scale; this adult growth rate can often be experimentally and/or observationally determined in a manner (relatively) independent from other process factors (see e.g. [8]). In addition, we choose the resulting characteristic seed dispersal distance as the characteristic length scale. For every model variable, the biomass density is scaled relative to the adult carrying capacity *Â*_max_. Furthermore, for algebraic convenience, all nondimensionalised model variables are divided by ratio of that variable’s growth rate relative to the characteristic growth rate *ĉ*_*A*_.

This leads to the following nondimensional reformulation of Equation (1)

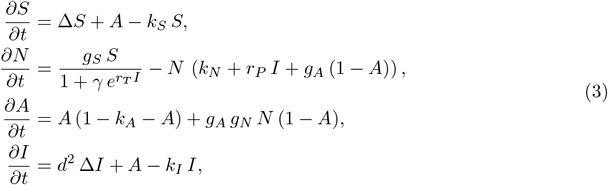

where 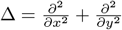 and the nondimensional parameters are given by

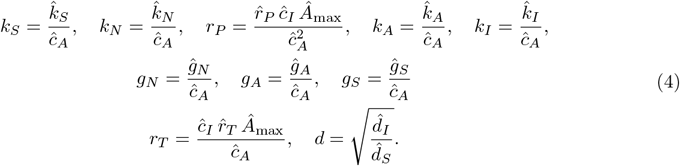

We note that, due to the range of ecological feasibility for our parameters reported in Table 1, we assume *k*_*S*_ *>* 0, *k*_*N*_ *>* 0, *r*_*P*_ ≥ 0, 0 *< k*_*A*_, *k*_*I*_ *>* 0, *g*_*N*_ *>* 0, *g*_*A*_ *>* 0, *r*_*T*_ ≥ 0, and *ε >* 0. Moreover, we assume that in the absence of seeds, seedlings and toxicity, the growth rate of adults is positive for all *Â >* 0, which implies

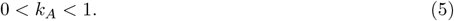

Ecologically feasible ranges of the nondimensionalised parameters, based on the associated dimensional values in Table 1, can be found in Table 2.

**Table 2:** Description and ranges of rescaled nondimensional parameters used in System (3), based on Table 1.

| Parameter | Description | Value |
| --- | --- | --- |
| $k_S$ | $S$ turnover rate | $1.3 - 2.0$ |
| $g_S$ | Growth rate of $S$ | $2.7 \cdot 10^{-7} - 1.3 \cdot 10^{-1}$ |
| $\gamma$ | Establishment sensitivity to toxicity parameter | $1.0 \cdot 10^{-5}$ |
| $r_T$ | Establishment sensitivity to toxicity parameter | $0 - 0.8 \cdot 10^4$ |
| $k_N$ | Death rate of $N$ | $0.8 \cdot 10^{-1} - 3.0$ |
| $r_P$ | Increased mortality of $N$ caused by $I$ | $0 - 1.0 \cdot 10^3$ |
| $g_A$ | Transition rate from $N$ to $A$ | $0.8 \cdot 10^{-3} - 4.0$ |
| $k_A$ | Mortality rate of $A$ | $4.0 \cdot 10^{-2}$ |
| $g_N$ | Transition rate from $S$ to $N$ | $1.0 - 1.0 \cdot 10^2$ |
| $k_I$ | Inhibitor decay rate | $2.8$ |
| $d$ | Square root of diffusion ratio | $0 - 1.8$ |

### Mathematical analysis: aims and goals

We determine the spatially homogeneous steady states (Section 3) and their linear stability with respect to spatially homogeneous and heterogeneous perturbations (Section 4). Furthermore, we investigate the presence of travelling waves (Section 5), determine properties of the wave profile, and determine the wave speed. To facilitate presentation, we organise the main results in Propositions and Theorems.

## 3. Spatially homogeneous steady states

For future reference and notational convenience, we introduce the establishment function

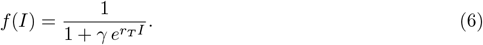

### Proposition 1

*System* (3) *admits two spatially homogeneous steady states*,

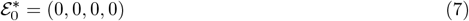

*and*

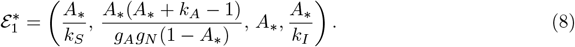

*Here, A*_*_ ∈ (1 − *k*_*A*_, 1) *is the unique solution to*

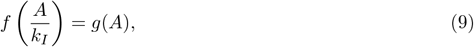

*where*

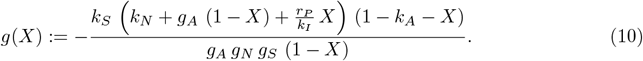

*Proof*. Spatially homogeneous steady states associated to system (3) are given by the solutions to

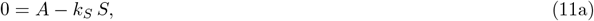

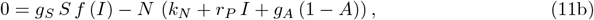

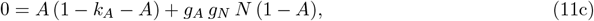

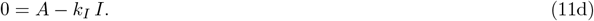

First, we observe that system (11) admits a trivial solution where all components vanish (representing bare soil):

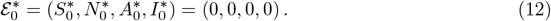

In order to compute the nontrivial equilibria of System (3), we first solve Equation (11a) and (11d) which lead to

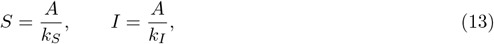

respectively. Substituting Equation (13) into Equation (11b) we obtain

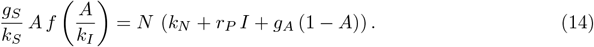

Solving Equation (14) for *N* we obtain

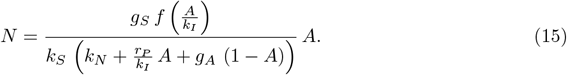

Substituting Equation (15) into Equation (11c) yields

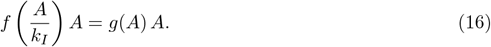

Clearly, *A* = 0 is a solution to (16), leading to the trivial solution 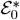 (12); division by *A* leads to (9).

It remains to show that Equation (9) has a unique nontrivial solution on the (ecologically feasible) interval (0, 1). We observe that for *X* ∈ [0, 1), the function *g*(*X*) satisfies the following properties:

- *g*(0) *<* 0,
- lim_*X→*1_ *g*(*X*) = +∞ (*g* has a vertical asymptote at *X* = 1),
- *g*″ (*X*) *>* 0 (*g* is convex),
- *g*(*X*) = 0 if and only if *X* = 1 − *k*_*A*_ (*g* has a unique root in the interval *X* ∈ (0, 1)).

Furthermore, the establishment function *f*(*X*) (6) satisfies the following properties:

- 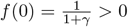
- *f*′ (*X*) *<* 0 for all *X* ∈ ℝ (*f* is strictly monotonically decreasing)
- *f*(*X*) *>* 0 for all *X* ∈ ℝ

Consequently, there exists a unique *A*_*_ ∈ (0, 1) that satisfies Equation 9. Moreover, since *f*(*X*) is positive and *g*(*X*) is positive only if *X >* 1 − *k*_*A*_, we find that *A*_*_ *>* 1 − *k*_*A*_; see also Figure 1. Therefore, we have that the unique nontrivial spatially homogeneous steady state of System (3) is given by

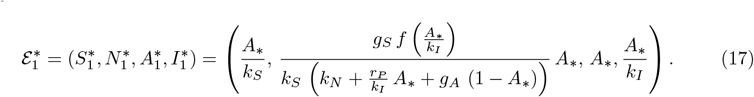

Using 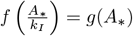 yields (8).

**Figure 1:**
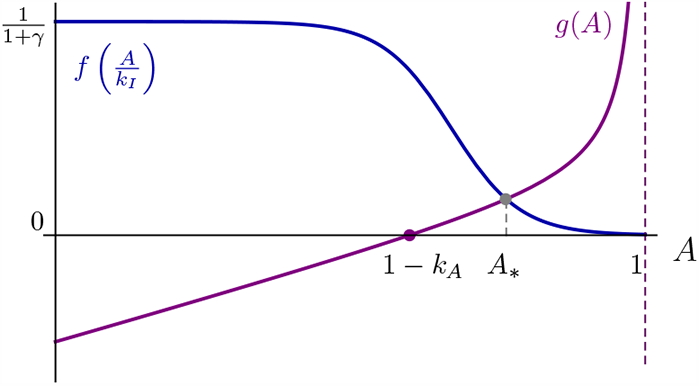
Schematic representation of the functions 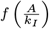 (blue solid line) and *g*(*A*) (purple solid line) as defined in Equation (6) and Equation (10), respectively, for *γ* = 10^*−*5^, *r*_*T*_ = 48, *k*_*S*_ = 1.3, *k*_*N*_ = 0.08, *g*_*A*_ = 1, *r*_*P*_ = 0, *k*_*I*_ = 2.8, *k*_*A*_ = 0.4, *g*_*N*_ = 13, and *g*_*S*_ = 0.13. The purple dot corresponds to the unique zero of *g*(*A*) in the interval *A* ∈ [0, 1], whereas the gray dot corresponds to the unique *A* ∗ where 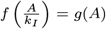 i.e. the *A*-component of the unique nontrivial steady state of System (3).

For future reference, based on the results of Proposition 1, we write

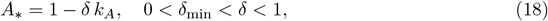

see also Figure 1. The lower bound *δ*_min_ can be determined by observing that the establishment function *f*(*I*) is bounded above by 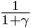 and hence 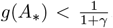. Solving 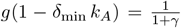 leads to

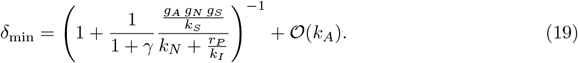

Moreover, according to the parameter values reported in Table 2, we have *k*_*A*_ = 0.04 ≪ 1. In the upcoming analysis, we will occasionally use *k*_*A*_ as a (regular) perturbation parameter, to gain insight into the solutions of complicated algebraic equations.

## 4. Linear stability

### 4.1. Spatially homogeneous perturbations

In this section, we analyse the linear stability of spatially homogeneous steady states 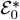(7) and 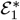 (8) with respect to spatially homogeneous perturbations.

#### Proposition 2

*The trivial steady state* 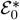 (7) *is unstable with respect to spatially homogeneous perturbations. The nontrivial steady state* 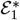 (8) *is linearly stable with respect to spatially homogeneous perturbations, as long as*

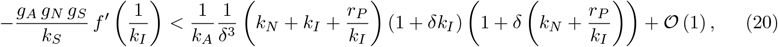

*assuming* 0 *< k*_*A*_ ≪ 1.

*Proof*. The Jacobian matrix corresponding to System (3) reads

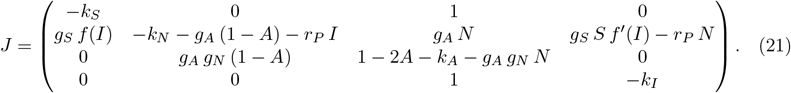

The characteristic polynomial associated to *J* evaluated at 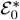 is given by

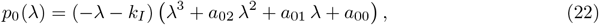

where

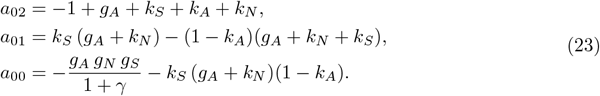

The polynomial *p*_0_(*λ*) admits four roots 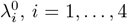. We identify 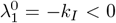, whereas the sign of the other three eigenvalues is investigated using the Routh-Hurwitz criterion. In particular, 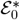 is asymptotically linearly stable if and only if the roots of the third order polynomial in (22) have negative real part, i.e. if and only if

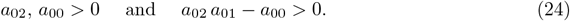

Due to the non-negativity constraints on our parameters and the bound on *k*_*A*_ (5), we have *a*_00_ *<* 0, thereby violating the Routh-Hurwitz criterion. Consequently, at least one of the three eigenvalues 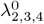 has positive real part, and the equilibrium 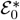 is unstable with respect to spatially homogeneous perturbations.

Concerning 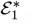 we define two new parameters *η, ζ >* 0 as

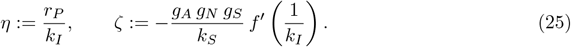

As long as *η, ζ* and all parameters in System (11) are O(1) with respect to *k*_*A*_ ≪ 1, the characteristic polynomial associated to *J* evaluated at 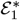 is given by

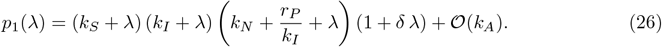

Due to the non-negativity assumption on the model parameters, *p*(*λ*) admits four negative roots, which in turn implies that 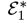 is a stable steady state w.r.t. homogeneous perturbations. The case where *η* and/or *ζ* are much larger than 𝒪(1) is analysed in Appendix Appendix A. The outcome of this analysis is that all eigenvalues have negative real part as long as 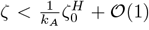, with the Hopf bifurcation threshold 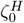 given by (A.3). Substituting *η* and *ζ* (25) yields (20).

#### Corollary 1

*The nontrivial steady state* 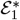 (8) *is stable with respect to spatially homogeneous perturbations for the parameter ranges in Table 2*.

*Proof*. From (6), we see that 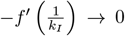 as *r*_*T*_ ↓ 0. Moreover,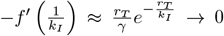 as *r*_*T*_ → ∞. The function 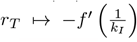 has a unique maximum, attained at 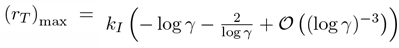, with value 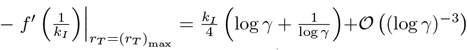.Implementing the values of Table 2, we combine the above with 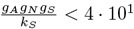 to obtain

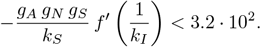

From the same Table, we infer

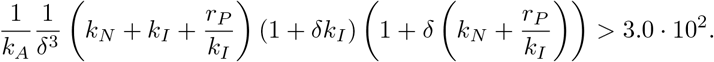

We see that straightforward estimates do not suffice to conclude that (20) is satisfied for the parameter ranges in Table 2; however, the bounds are sufficiently close to conclude that the region in parameter space for which 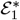 is unstable with respect to spatially homogeneous perturbations is relatively small. Furthermore, the value of *δ* is determined by the other system parameters through (9) and (18). Hence, we numerically determine the maximal real part of the eigenvalues of 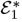, by first determining the value of *A*_*_ (cf. Proposition 1) and then calculating the eigenvalues of the associated Jacobian. For the parameter ranges in Table 2, the maximum real part of the eigenvalues is found to be − 0.96 *<* 0. Hence, 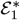 is stable with respect to spatially homogeneous perturbations for the parameter ranges in Table 2.

### 4.2. Spatially heterogeneous perturbations

Since Turing patterns can emerge when steady states are stable with respect to spatially homogeneous perturbations but lose their stability when considering spatially heterogeneous perturbations, in this section we focus our attention only on the steady state 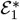 (see Section 4.1).

#### Proposition 3

*The spatially homogeneous steady state* 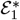 (8) *is linearly stable with respect to spatially heterogeneous perturbations, as long as*

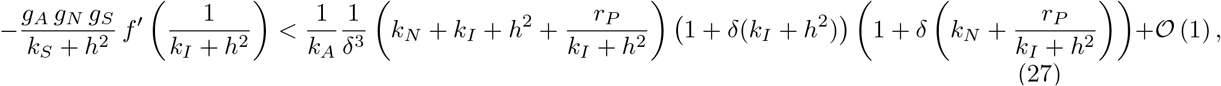

*for all h* ∈ ℝ, *assuming* 0 *< k*_*A*_ ≪ 1.

*Proof*. We introduce the following non-uniform perturbations:

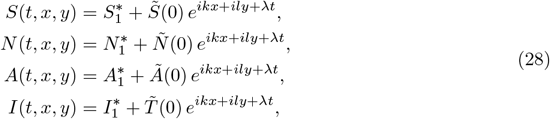

where the (spatial) wave number of the perturbation is defined as 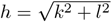 and *λ* represents the temporal growth. Linearising System (3) around 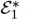, we obtain the following system for the perturbations 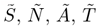 defined in (28):

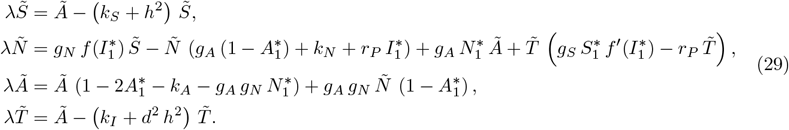

System (29) can be written as an eigenvalue problem 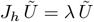, where 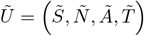 and

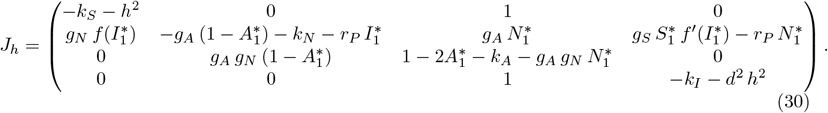

We observe that this eigenvalue problem can be made identical to the eigenvalue problem for spatially homogeneous perturbations, as studied in the proof of Proposition 2, by replacing

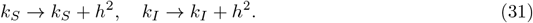

Hence, the same stability criterion as in Proposition 2 applies, with the substitution (31).

#### Corollary 2

*The nontrivial steady state* 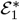(8) *cannot undergo a Turing bifurcation for the parameter ranges in Table 2*.

*Proof*. From the proof of Proposition 2, we see that −*f*′(*X*) has a unique maximum at a fixed value of 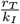 for *γ* fixed. The right hand side of (27) is by construction independent of *r*_*T*_, since *r*_*T*_ only occurs in the derivative of *f* (encoded by *ζ*), and the right hand side of (27) is an expansion of a bound on *ζ*.

From Proposition 2, we know that 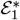 is stable with respect to spatially homogeneous perturbations, which is equivalent to setting *h* = 0 in (27). We infer from Proposition 2 that (27) is satisfied for *h* = 0 for all admissible parameter ranges in Table 2. Therefore, for given *k*_*I*_ and *γ*, we may assume without loss of generality that *r*_*T*_ is chosen such that 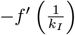 is maximal, as (*r*_*T*_)_max_ = 3.3 · 10^1^ falls (well) within the admissible range of *r*_*T*_. When *k*_*I*_ → *k*_*I*_ + *h*^2^ is increased, this argument continues to hold until *r*_*T*_ reaches its maximal admissible value; when *h* is increased beyond this point, 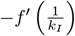 is smaller than its unique maximal value. Combining this argument with 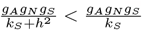, we see that

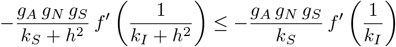

for all *h* ∈ ℝ.

The right hand side of (27) is non-monotonic in *h*. However, a lower bound is found analogously to the proof of Corollary 1 by setting *r*_*P*_ = 0 and minimising the other parameters. Since all components of the right hand side of (27) are increasing functions of *h*^2^, the value of the right hand side of (27) is bounded from below by its value for *h* = 0. The above arguments imply that (27) therefore remains satisfied for *h >* 0.

We conclude that no Turing(-Hopf) bifurcation can take place for the parameter ranges in Table 2.

## 5. Travelling waves

As shown in Section 4, for the parameter ranges in Table 2, the nontrivial steady state 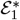 is spectrally stable (Corollary 1), whereas the trivial steady state 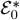 is unstable (Proposition 2). Moreover, numerical simulations of System (1) (on a sufficiently large, one-dimensional domain, with Neumann boundary conditions) show the emergence of travelling wave solutions invading the unstable steady state 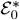 for a broad range of parameter values (see an example in Figure 2). These simulations suggest the existence of a travelling wave with fixed wave speed in System (1) on an unbounded one-dimensional spatial domain. In this section, we investigate the existence of such a travelling wave, and provide arguments for its existence in a large part of parameter space. Moreover, we show that the numerically measured wave speed coincides with the so-called linear spreading speed, to a high degree of accuracy. This suggests that the numerically observed front can be classified as a *pulled front*, that is, where the linear spreading of small perturbations pulls the front into the linearly unstable bare soil steady state [36].

**Figure 2:**
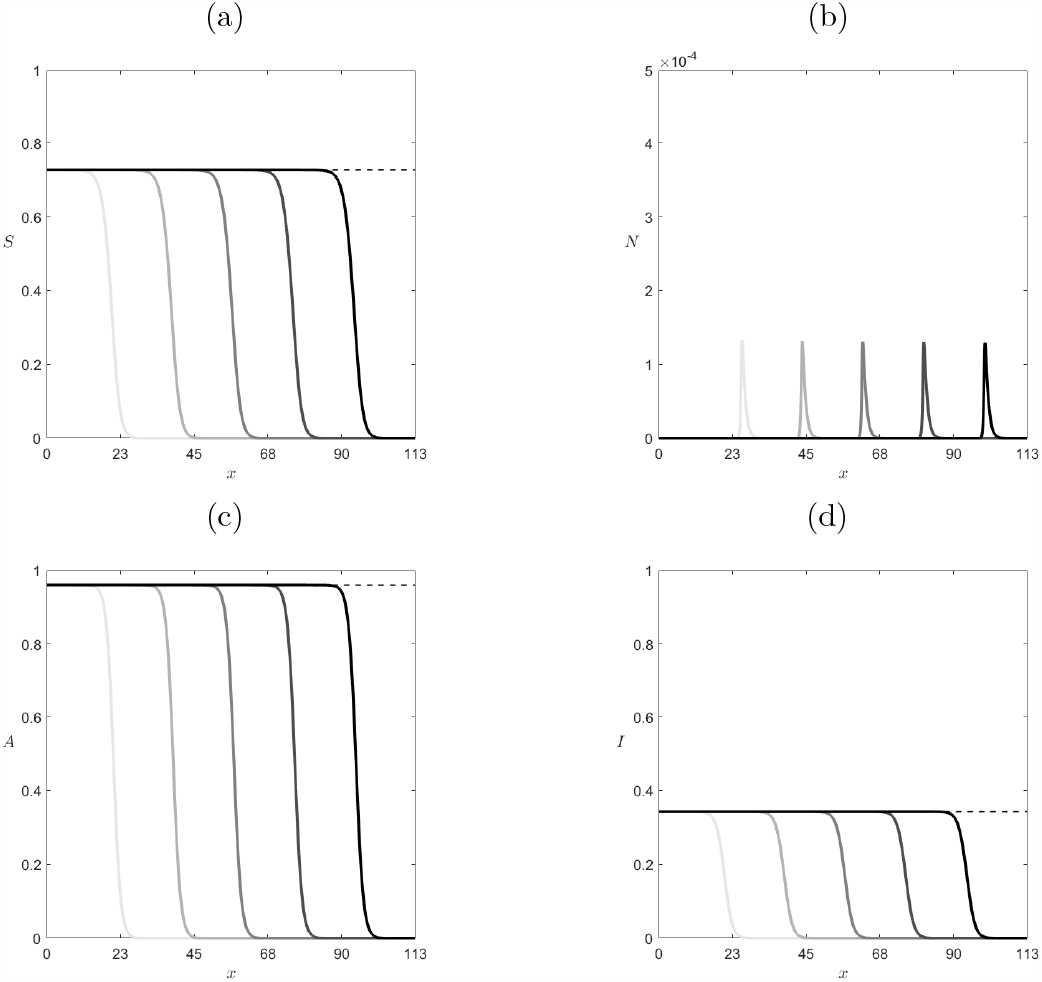
Numerical profiles for (a) *S*, (b) *N*, (c) *A*, and (d) *I* obtained by simulating Equation (3) for *t* ∈ [0, 87.5]. Profiles are shown a *t* distance Δ*t* = 17.5 for *g*_*S*_ = 0.132, *k*_*S*_ = 1.32, *g*_*N*_ = 20, *r*_*T*_ = 4080, *k*_*N*_ = 2, *r*_*P*_ = 480, *g*_*A*_ = 0.8, *d* = 0.913, and other parameters values as in Table 2. The intensity of the shading (from light gray to black) increases with *t*.

To prepare the analysis, we introduce a co-moving frame via the variable *ξ* = *x* − *c t*, where *c* represents the wave speed. System (3) hence becomes

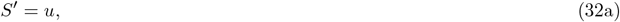

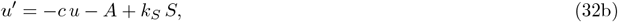

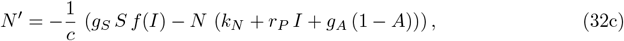

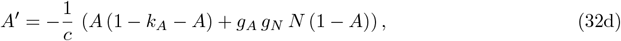

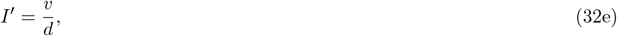

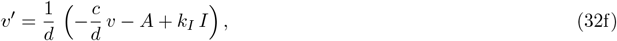

which can also be expressed in the compact form *z*′ = *F*(*z*), where *z* = (*S, u, N, A, I, v*). System (32) admits the two equilibria

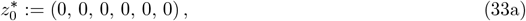

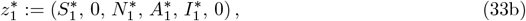

where the components of 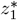 coincide with those defined in Equation (8). The equilibria 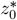 and 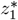 are the representation of the spatially homogeneous steady states 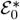 (7) and 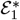 (8) in the travelling wave framework.

In this context, a right-moving front (with *c >* 0) invading the trivial steady state 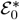 coincides with an heteroclinic connection from 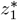 to 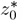. Such an orbit must therefore lie in the intersection of the unstable manifold of 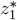 (denoted by 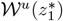 and the stable manifold of 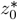 (denoted by 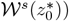. To investigate the potential existence of right-moving fronts we hence need to derive the parametric conditions such that

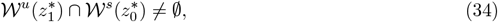

which follow directly from the investigation of the dimensions of the stable and unstable manifolds of 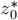 and 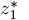.

The main point of our analysis consists in studying the characteristic polynomial associated to System (32), which can be expressed as

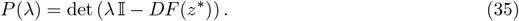

The roots of *P*(*λ*) evaluated at 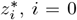 will provide information about the dimension of the stable and unstable manifolds of these equilibria, indicating whether Equation (34) can hold.

### 5.1. Local analysis of 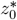

#### Theorem 1

*The dimensions of the stable and unstable subspaces of* 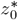 *(denoted as* 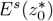 *and* 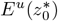, *respectively) satisfy*

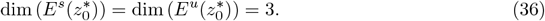

*Proof*. The characteristic polynomial of System (32) at 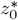 is given by

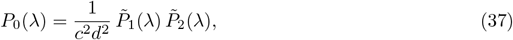

with

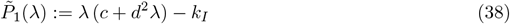

and

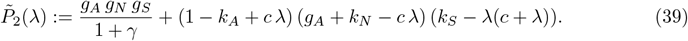

The quadratic function 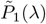 is convex and negative at *λ* = 0; therefore it admits two real roots of opposite sign, namely 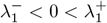. To study the roots of 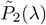, we write

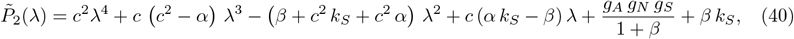

where *α* = *g*_*A*_ + *k*_*A*_ + *k*_*N*_ − 1 and *β* = (*g*_*A*_ + *k*_*N*_) (1 − *k*_*A*_); note that *β >* 0.

The sign of the roots of 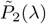 can be investigated applying the Routh-Hurwitz criterion, by rewriting Equation (40) as

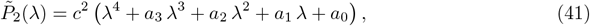

where

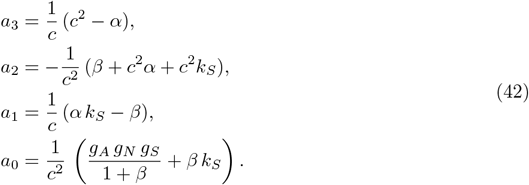

Applying Descartes’ rule of signs, which states that the number of roots with negative (resp. positive) real part corresponds to the number of sign changes (resp. permanences) on the coefficients of 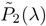, and taking into account the fact that *a*_0_ *>* 0, we observe that the conditions *a*_3_ *>* 0, *a*_2_ *>* 0, and *a*_1_ *>* 0 cannot be verified simultaneously, i.e. there is at least one sign variation and one permanence. Hence, the fourth order polynomial 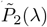 admits at least one root with positive and one with negative real part, denoted by 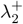 and 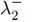.

Moreover, there are no purely imaginary roots of 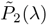 since 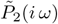 is a real polynomial if and only if 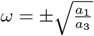 and *a*_1_, *a*_3_ have the same sign only for *a*_2_ *<* 0, which implies

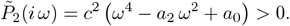

Consequently, we have (considering 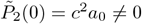) that the centre eigenspace 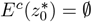, from which it follows that the phase space can be decomposed into the direct sum of the stable and unstable eigenspaces 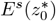 and 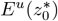 respectively, i.e. 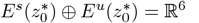.

Besides the two roots with opposite real signs 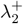 and 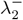 derived above, we need to check the sign of the other two roots of 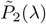, which we define as 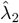 and 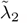. We analyse all possible scenarios:

- If 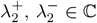, then 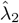 and 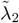 must be equal to the complex conjugates of 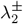, i.e. 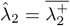 and 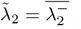.
- If 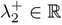, then 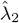 must be positive and real. This is due to the fact that 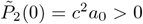 and 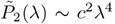 as *λ* → ∞; therefore, the graph of 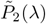 must have an even number of crossings with the positive horizontal axis.
- Analogously, if 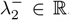, then 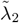 must be negative and real. This is due to the fact that 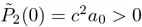 and 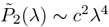 as *λ* → −∞; therefore, the graph of 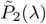 must have an even amount of crossings with the negative real axis.

To summarize, we conclude that *P*_0_(*λ*) admits a total of three eigenvalues with positive real part (namely 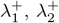 and 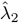), and three with negative real part (namely 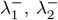 and 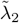), which leads to the claim of the theorem.

### 5.2. Local analysis of 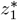

#### Theorem 2

*Assume* 0 *< k*_*A*_ ≪ 1 *is sufficiently small. For* 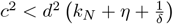, *the dimensions of the stable and unstable eigenspace of* 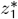 *(corresponding to* 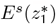 *and* 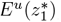, *respectively) satisfy* dim 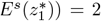 *and* dim 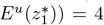. *Only when* 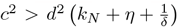 *there exists a* 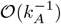*-value of ζ* = *ζ*_*H*_ *such that, if ζ > ζ*_*H*_, dim 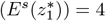 *and* dim 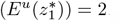.

*Proof*. The characteristic polynomial of System (32) at 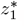, in agreement with Equation (35), is defined as

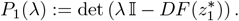

Our analysis is based on the observation made at the end of Section 3 that, since *k*_*A*_ ≪ 1, we can treat *k*_*A*_ as an asymptotically small perturbation parameter to investigate the roots of complicated algebraic expressions such as (35).

We write *A*_*_ = 1 − *δ k*_*A*_ with 0 *< δ <* 1, cf. (18). Recalling the definition of *η* and *ζ* in (25), we expand *P*_1_(*λ*) for small *k*_*A*_ and consider the four regimes

I. *η, ζ* ∈ 𝒪(1),
II. *η* ≫ 1, ζ ∈ 𝒪(1),
III. *η* ∈ 𝒪(1), *ζ* ≫ 1,
IV. *η, ζ* ≫ 1.

#### Regime I

*η, ζ* ∈ 𝒪(1).. In this regime, the characteristic polynomial can be expressed as

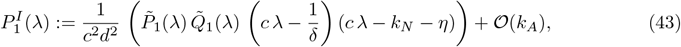

where

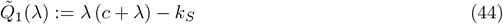

and 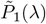 as defined in (38). In the proof of Theorem 2, it is shown that the roots of 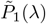 are real and have opposite sign, i.e. 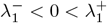. The same statement holds for the roots of 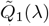, since 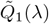 is convex and 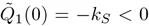; we denote the roots of 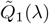 as 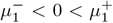. The two remaining roots of the leading order expression of 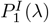 (43) are given by 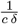 and 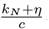, which are both real and positive. All roots of the leading order expression of 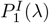 are nondegenerate and bounded away from zero, and therefore perturb regularly for *k*_*A*_ ≪ 1. Therefore, dim 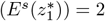 and dim 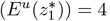.

#### Regime II

*η* ≫ 1, *ζ* ∈ 𝒪(1).. The 𝒪(*k*_*A*_) terms in the expansion of 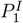 (43) do not depend on *η*. Hence, the roots of *P*_1_ in regime II are equal to those in regime I. The only difference is that now the eigenvalue 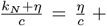 𝒪(1), but this does not affect its sign, which remains positive.

Therefore, as in Regime I, we obtain dim 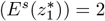 and dim 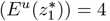.

#### Regime III

*η* ∈ 𝒪(1), *ζ* ≫ 1.. In this regime, the characteristic polynomial is to leading order given by

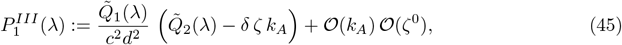

where

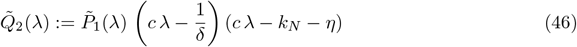

and 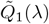 as defined in (44). We therefore need to split our investigation into further subcases depending on the magnitude of *ζ k*_*A*_.

i. *ζ k*_*A*_ ≪ 1: here 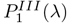 is a regular perturbation of 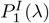; we obtain same result on the sign of the eigenvalues, that is, dim 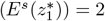 and dim 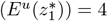.
ii. *ζ k*_*A*_ = 𝒪(1): we write 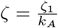 with *ζ*_1_ ≥ 0. Substituting this assumption into Equation (45) leads to

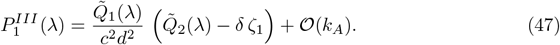

When *ζ*_1_ = 0, we have that 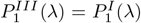, hence the sign of the eigenvalues is identical. When *ζ*_1_ *>* 0, since 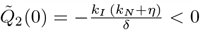 and 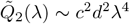 as *λ* → ±∞, we have that 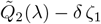 must admit at least one positive and one negative real root. For *ζ*_1_ sufficiently small, the roots of 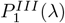 are a regular perturbation of the roots of 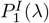, which implies that the two remaining roots of 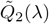 are real and positive. As *ζ*_1_ is increased, this root pair undergoes a (stabilising) Hopf bifurcation for sufficiently large values of *ζ*_1_, namely for 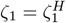 where

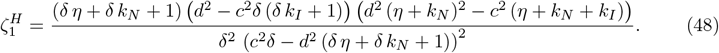

In other words, the real part of the complex conjugate roots of 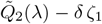 is positive for 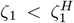, vanishes for 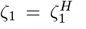, and is negative for 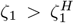. The expression for 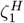 in (48) is derived by solving 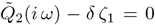 and imposing that the imaginary part of the resulting polynomial is zero. This gives an expression for *ω* which can be substituted back into 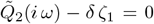; then we can subsequently solve this equation for *ζ*_1_ to obtain 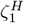. Imposing the feasibility conditions *ω*^2^ *>* 0 and *ζ*_1_ *>* 0 we obtain that a Hopf bifurcation occurs if and only if

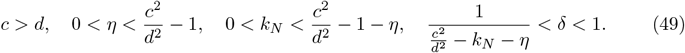

In particular, the above conditions hold if and only if 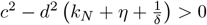.
iii. *ζ k*_*A*_ ≫ 1: In this case, the equation 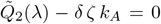 implies that |*λ*| ≫ 1. To leading order, we thus have

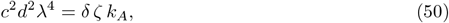

which is solved by two real and two purely complex roots, namely

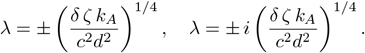

Since the complex roots are purely imaginary to leading order, these need further unfolding to determine the sign of their real part. Including higher order terms (𝒪(*λ*^3^) and 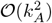, respectively) in Equation (50) leads to the following refinement of the complex roots

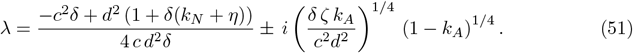

We have two possibilities:
  - If 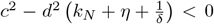, the real part of the roots in (51) is positive. Therefore, taking into account the sign of the other roots of *P*_*III*_ (*λ*), we find dim 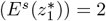 and dim 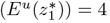.
  - If 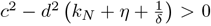, the real part of the roots in (51) is negative, and we find dim 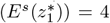 and dim 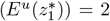. Note that, comparing to the case *ζk*_*A*_ ≪ 1, this implies that somewhere between *ζk*_*A*_ ≪ 1 and *ζk*_*A*_ ≫ 1, a sign change must have occurred. This is precisely the Hopf bifurcation found at *ζk*_*A*_ = 𝒪(1), to wit, at 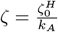 (48).

#### Regime IV

*η, ζ* ≫ 1.. To leading order, the characteristic polynomial in this regime coincides with the one in Regime III, i.e.

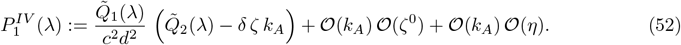

As in Regime III, we need to consider different relations between the orders of *η* and *ζ k*_*A*_ to determine the roots of 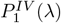.

i. *η* ≫ *ζ k*_*A*_: In this case, 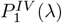 is a regular perturbation of 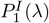; its roots are then given by 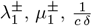, and 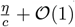.
ii. *η* ∼ *ζ k*_*A*_: Here we can express 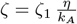, with *ζ*_0_ = 𝒪(1). To leading order, we obtain

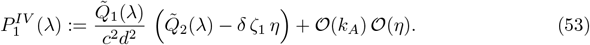

The roots of (53) are studied by distinguishing two regimes. Focusing on |*λ*| ≫ 1; in this case, 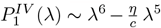, we obtain that one root is, to leading order, given by 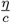. The other five roots are studied by investigating 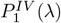 to leading order in *η* for *λ* = 𝒪(1), i.e.

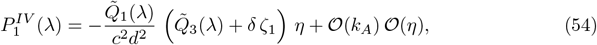

where

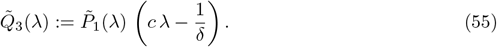

Two roots are to leading order given by the roots of 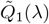. For the other three, we see that, since 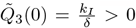 and 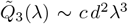 as *λ* → ±∞, the polynomial 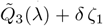 always admits at least one negative real root. As for the other two, we observe that no Hopf bifurcation occurs in this case (since the only solution to 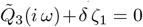 is given by *ω* = 0). The sign of their real part hence remains the same as *ζ*_1_ is varied, and since we know that for *ζ*_1_ = 0 the other two roots of 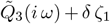 are real and positive, they remain positive for all *ζ*_1_.
iii. *η* ≪ *ζ k*_*A*_: In this case, the characteristic polynomial is given by

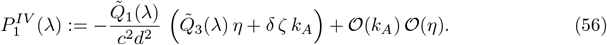

Two roots are to leading order given by the roots of 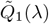. Solving 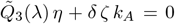 hence requires |*λ*| ≫ 1. Expanding for large *λ* yields

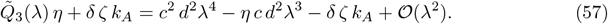

Note that, as − *δ ζ k*_*A*_ *<* 0 and *c*^2^ *d*^2^*λ*^4^ − *η c d*^2^*λ*^3^ − *δ ζ k*_*A*_ ∼ *c*^2^*d*^2^*λ*^4^ as *λ* →±∞, the leading order polynomial (57) has at least two real roots of opposite sign. To further determine the roots of (57), we consider four possible balances:
  - If *c*^2^ *d*^2^*λ*^4^ ∼ *η c d*^2^*λ*^3^ ≫ *δ ζ k*_*A*_, Equation (57) reduces to leading order to *c*^2^ *d*^2^*λ*^4^ − *η c d*^2^*λ*^3^ = 0; this equation admits one real positive root λ = 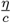 and one zero root with multiplicity three, that needs further unfolding. For *λ* ∼ 0, Equation (57) admits the three roots

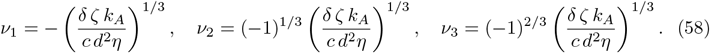

The roots *ν*_1_ and *ν*_3_ have negative real part, whereas *ν*_2_ has positive real part.
  - If *η c d*^2^*λ*^3^ ∼ *δ ζ k*_*A*_ ≫ *c*^2^ *d*^2^*λ*^4^, to leading order Equation (57) admits the roots in (58), whose sign has been analysed in the previous balance point. The fourth root of (57) is real and negative.
  - If *c*^2^ *d*^2^*λ*^4^ ∼ *δ ζ k*_*A*_ ≫ *η c d*^2^*λ*^3^, we have two real and two purely complex roots, namely

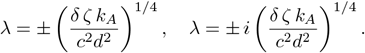

The sign of the real part of the complex roots is obtained by considering the higher order term *η c d*^2^*λ*^3^, from which we get

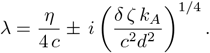

In this case we hence have one root with negative real part and three roots with positive real part.
  - If *c*^2^ *d*^2^*λ*^4^ ∼ *η c d*^2^*λ*^3^ ∼ *δ ζ k*_*A*_, we can define *λ* := *λ*_0_ *η* and *ζ k*_*A*_ := *ζ*_2_ *η*^4^. Equation (57) thus becomes

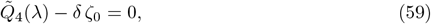

where 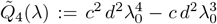. This function satisfies 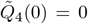 and 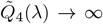 as *λ* → ±∞, therefore Equation (59) admits two real roots of opposite sign for any *ζ_2_ >* 0. The two other complex roots have positive real part equal 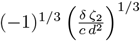 to leading order; since no Hopf bifurcations are possible, the sign of the real part of the complex roots remains positive for any *ζ*_2_ *>* 0. Hence also in this case we have one root with negative real part and three roots with positive real part.

Consequently, considering all possible balances we find for Regime IV that dim 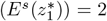 and dim 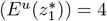.

#### Conclusion

We observe that 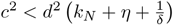 is automatically satisfied when *η* ≪ 1, that is, in Regime II and Regime IV. Combining the results from Regimes I–IV, we see that dim 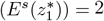 and dim 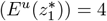 when 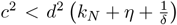. Only when 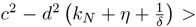 0 there exists a 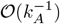-value of *ζ* = *ζ*_*H*_ such that, if *ζ > ζ*_*H*_, dim 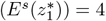 and dim 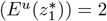.

### 5.3. Existence of a travelling wave

In phase space, a travelling wave solution corresponds to a heteroclinic orbit connecting 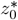 and 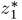, thus lying in the intersection of the unstable manifold of one equilibrium and the stable manifold of the other. We use the Local Stable Manifold Theorem to infer from Theorem 1 that the dimensions of the stable and unstable manifolds of 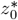 are

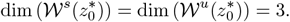

Likewise, we infer from Theorem 2 that the dimensions of the stable and unstable manifolds of 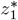 are either

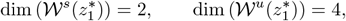

provided 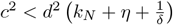, or

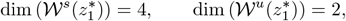

provided 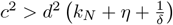 and *ζ* (25) is sufficiently large, in particular 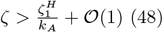.

Recall that the aim of this section is to obtain analytical insight into numerically observed travelling fronts that invade the trivial steady state, which for a right-moving front with positive speed *c* corresponds to a heteroclinic connection *from* 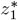 *to* 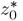. Therefore, we take 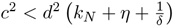, for reasons to be explained momentarily. Observing that codim 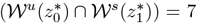, whereas codim 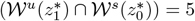, and taking into account the fact that the phase space is six-dimensional, we conclude that generically 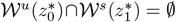 and dim 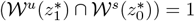 (if non-empty). In the latter case, this intersection is generically transversal and hence persists when *c* is perturbed. This leads us to the following Corollary:

#### Corollary 3

*We generically expect a one-parameter family of heteroclinic connections from* 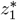 *to* 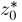, *parametrised by the wave speed c, with* 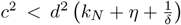. *Moreover, we expect this family to exist in an open region of parameter space. Every member of this family corresponds to a right-moving front invading the trivial steady state* 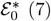.

Note that the above arguments do not constitute a proof of the existence of a heteroclinic connection from 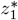 to 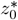, as the intersection 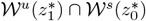 might be empty. However, in the upcoming section, we identify a parameter range for which such a heteroclinic connection exists (Theorem 3).

#### Remark 1

*The same reasoning can be applied to generically expect the existence of a heteroclinic connection from* 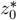 *to* 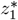, *for sufficiently large wave speeds* 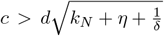 *and sufficiently large values of ζ. However, for all c* 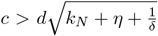, *we have that* 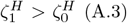, *and ζ does not exceed* 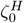 *for the parameter ranges in Table 2, cf. Corollary 1. For this reason, we do not investigate this anomalous wave any further in the current paper*.

### 5.4. Properties of the wave profile

In this section, we derive generic properties satisfied by a right-moving travelling front solution to System (3), which is equivalent to a heteroclinic connection from 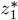 to 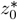 in System (32). These properties can be used to explore the connection between such a travelling wave and the Janzen-Connell distribution.

#### Lemma 1

(Monotonicity of *S* and *I*). *Let* (*S*(*ξ*), *u*(*ξ*), *N* (*ξ*), *A*(*ξ*), *I*(*ξ*), *v*(*ξ*)) *be a solution to System* (32) *representing a right-moving front travelling with speed c >* 0. *If A*′ (*ξ*) *<* 0 *for all ξ, then S*^*I*^(*ξ*) *<* 0 *and I*′ (*ξ*) *<* 0 *for all ξ*.

*Proof*. We first consider *S*′ (*ξ*). The proof strategy is based on deriving an explicit solution for *S*(*ξ*) by means of a Green’s function, which in turn allows us to obtain an explicit solution for *S*′ (*ξ*) as a function of *A*′ (*ξ*) using integration by parts.

We write equations (32a)-(32b) as a single second order equation for *S*, yielding

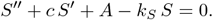

The boundary conditions

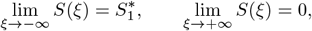

uniquely determine the solution

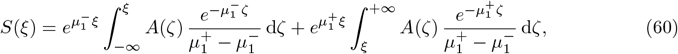

with

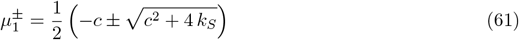

the roots of 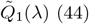. Consequently, we have that

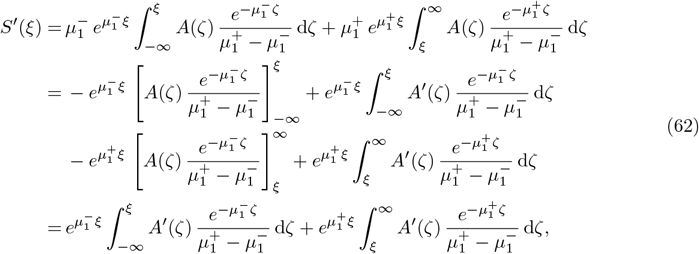

which is negative if *A*′ (*ξ*) *<* 0 for all *ξ* ∈ ℝ. The proof of *I*′ (*ξ*) *<* 0 is analogous, with 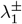, the roots of 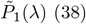 (38), replacing 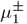.

#### Lemma 2

(Monotonicity of *A*). *Let* (*S*(*ξ*), *u*(*ξ*), *N* (*ξ*), *A*(*ξ*), *I*(*ξ*), *v*(*ξ*)) *be a solution to System representing a right-moving front travelling with speed c >* 0. *Then A*(*ξ*) *<* 1 *for all ξ. Moreover, there exists a ξ*_0_ ∈ ℝ *such that*

- *A*(*ξ*) *>* 1 − *k*_*A*_ *for all ξ < ξ*_0_, *and*
- *A*(*ξ*) *<* 1 − *k*_*A*_ *and A*^*I*^(*ξ*) *<* 0 *for all ξ* ≥ *ξ*_0_.

*Proof*. From Equation (32d) together with the positivity assumptions on *N* and the parameters *g*_*A*_, *g*_*N*_, it follows that when *A* ≥ 1, then *A*′ *>* 0. Therefore, if solution crosses the threshold *A* = 1 for a certain *ξ* = *ξ*_1_, it will remain above *A* = 1 for all *ξ > ξ*_1_. This contradicts the assumption on the travelling wave solution, that *A*(*ξ*) → 0 as *ξ* → ∞.

Again from Equation (32d) together with the positivity assumptions on *N* and the parameters *g*_*A*_, *g*_*N*_, it follows that when *A* ≤ 1 − *k*_*A*_, then *A*′ *<* 0. Therefore, if solution crosses the threshold *A* = 1 − *k*_*A*_ for a certain *ξ* = *ξ*_0_, it will remain below *A* = 1 − *k*_*A*_ for all *ξ > ξ*_0_. Since the travelling wave solution has *A*(*ξ*) → 0 as *ξ* → ∞ and *A*(*ξ*) → *A*_*_ *>* 1 − *k*_*A*_ as *ξ* → −∞, it follows that the solution crosses the threshold 1 − *k*_*A*_ for some *ξ*_0_. Once the solution has crossed this threshold, it will continue to decrease (strictly monotonically) to zero.

#### Remark 2

*When* 1 − *k*_*A*_ *< A <* 1, *no monotonicity of A is generically guaranteed. By redefining A* = 1 − *k*_*A*_ *a and linearising System* (32) *around the nontrivial equilibrium A*_*_ *(corresponding to a* = *δ), we have seen in Section 5.2 that in our travelling wave framework there are four unstable eigenvalues, hence leading to four associated eigenvectors with a nonzero a-component as follows:*

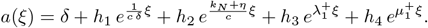

*Therefore, depending on the signs of the constants h*_*i*_, *i* = 1, …, 4, *A can admit several local minima and maxima in a neighbourhood of A*_*_.

Lemma’s 1 and 2 provide information on the monotonicity of *S, I* and *A*. However, for the seedling component *N*, one cannot derive monotonicity properties in full generality, due to the nature of the nonlinearity of Equation (32c).

To mitigate this problem, we consider the establishment function *f*(*I*) (6) for large values of *r*_*T*_. We observe that, for sufficiently large *r*_*T*_, *f*(*I*) behaves like a switch function (see also Figure 3):

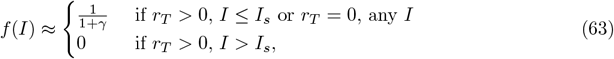

where *I*_*s*_ corresponds to the inflection point of *f* given by

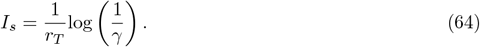

#### Theorem 3

(JC for strong toxicity and slow seed growth). *Let r*_*T*_ *be sufficiently large and g*_*S*_ *sufficiently small. Then, there exists a heteroclinic orbit in* (32) *from* 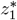 *to* 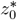 *for which the N-profile has a unique maximum, and the S-, A- and I-profiles are strictly monotonic*.

*Proof*. For asymptotically large *r*_*T*_, the establishment function *f*(*I*) (6) is exponentially close to 0 for *I > I*_*s*_ + *I*_1_ and exponentially close to 1 for *I < I*_*s*_ − *I*_1_, with *I*_1_ = (*r*_*T*_)^*α−*1^, for any 0 *< α <* 1. Moreover, both *I*_*s*_ (64) and *I*_1_ are asymptotically close to zero.

**Figure 3:**
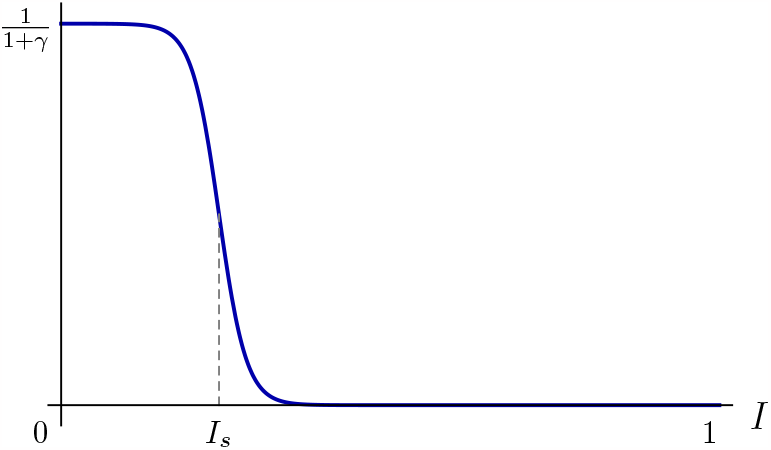
Schematic representation of the establishment function *f*(*I*) as defined in Equation (6), for large values of *r*_*T*_.

For *f*(*I*) ≡ 0, the hyperplane {*N* = 0} is invariant under the flow of (32). Moreover, for this choice of *f*, the nontrivial equilibrium 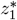 lies on {*N* = 0}, and is given by 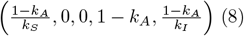. By Lemma’s 1 and 2, we see that *S, I* and *A* are strictly monotonically decreasing on the invariant hyperplane {*N* = 0}. Since {*N* = 0} is normally hyperbolic and *f*(*I*) is asymptotically small for all *I > I*_*s*_ + *I*_1_, the half-hyperplane *P*_0_ := *N* = 0, *I > I*_*s*_ + *I*_1_ perturbs to a locally invariant codimension-1 manifold *P* for the full system (32) [17]. The unstable manifold of 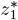 of the flow on *P*, that we denote by 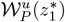, is 3-dimensional. Since *P* is normally repelling in the *N* -direction, we can conclude that in a neighbourhood of *P*, the 4-dimensional unstable manifold of 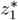 in the full system (32) is foliated as 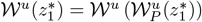.

The *A*-dynamics on *P* are to exponential order in given by 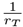 given by 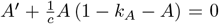, which yield 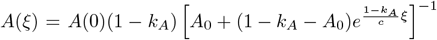. From the (linear) *S*- and *I*-dynamics on *P*, which depend linearly on *A*, we see that if 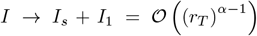, then both 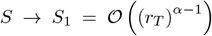 and 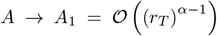; the same holds for the derivatives *u* and *v*. Moreover, in a sufficiently small neighbourhood of *P*, the normal *N* -dynamics are to leading order linear, and 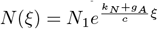 for *N*_1_ sufficiently small.

We investigate the intersection of 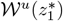 and 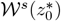 in a neighbourhood of *P*, and in a neighbourhood of *I* = *I*_*s*_. Close to both *P* and *I*_*s*_, the *S*-, *u*-, *A*-, *I*-, and *v*-components of orbits in 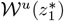 are 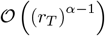, while *N* is sufficiently small by assumption. Hence, in order for 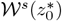 to intersect 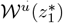 in this neighbourhood, all components of 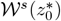 must be close to zero. It follows that the if intersection of 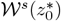 and 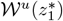 lies close to *P* and *I*_*s*_, it has to be close to the origin 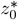. Close to the origin, the dynamics on 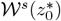 are linear, and 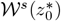 is close to 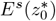. Hence, for *r*_*T*_ sufficiently large, transversal intersections of 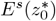 and 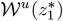 perturb regularly to transversal intersections of 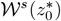 and 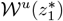.

Now, let *g*_*S*_ ≪ 1. For *g*_*S*_ = 0, the flow of (32) is equal to the flow of (32) under the assumption *f*(*I*) ≡ 0. Hence, the hyperplane {*N* = 0} is invariant when *g*_*S*_ = 0. Moreover, the trivial equilibrium 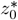 lies on {*N* = 0}. Solving the equations for *A, S* and *I* on {*N* = 0} yields the following unique heteroclinic orbit from 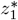 to 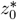 on {*N* = 0}:

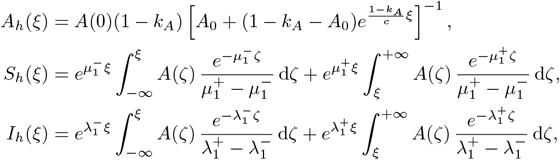

cf. Lemma 1. Thus, for *g*_*S*_, 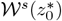 and 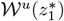 intersect transversally in the hyperplane {*N* = 0}, and this intersection is one-dimensional.

We investigate how this intersection perturbs for 0 *< g*_*S*_ ≪ 1. As the term *g*_*S*_ *S f* (*I*) is a regular perturbation of system (32), we know that both the hyperplane {*N* = 0} and the stable/unstable manifolds of the equilibria 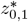 perturb regularly in *g*_*S*_. To determine the *N* -profile, we consider the unstable eigenvalues 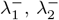 and 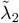 of 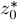 (cf. the proof of Theorem 1), which for 0 *< g*_*s*_ ≪ 1 can be determined explicitly: we find 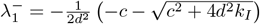 (38),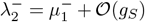 (61) and 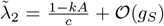, cf. (**??**)-(39). The associated eigenvectors can be determined by an expansion in powers of *g*_*S*_. To leading order, the *N* -component of the eigenvectors is zero, due to the fact that the intersection of 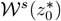 and 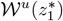 lies in the *N* = 0 hyperplane for *g*_*S*_ = 0. The first order correction of the eigenvectors associated to 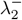 and 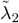 yields a nonzero *N* -component, to wit 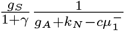 for the 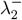 eigenvector and 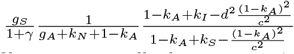 for the 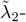 eigenvector. Hence, to first order in *g*_*S*_, the *N* -profile is exponentially decreasing as *ξ* → ∞, along these eigenvectors.

Now, define *ξ*_*_ through *I*_*h*_(*ξ*_*_) = *I*_*s*_ (64). Note that *ξ*_*_ is well-defined since *I*_*h*_ is strictly monotonically decreasing when *I*_*h*_ is sufficiently small (Lemma 1). *I* is to exponential accuracy in 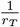 approximated by *I*_*h*_ on *P*, (at least) up to *I* = *I*_*s*_ + *I*_1_. In addition, *I* is to leading order in *g*_*S*_ approximated by the linear dynamics on 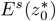, (at least) up to *I*_*s*_ − *I*_1_. The change of *I* over the interval (*I*_*s*_ − *I*_1_, *I*_*s*_ + *I*_1_) is small, and since system (32) is regularly perturbed for small *g*_*S*_ and large *r*_*T*_, this implies that the change in all other components over the *ξ*-interval associated to the change from *I* = *I*_*s*_ + *I*_1_ to *I* = *I*_*s*_ − *I*_1_ is small as well. Hence, to leading order, we can match the linear dynamics on 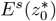 to the dynamics of 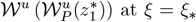. The transversality of the intersection of 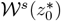 and 𝒲*u*(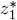 ensures that this matching procedure can be carried out for every component. The result for the *N* -profile is a single peak, to leading order in *g*_*S*_ up to *ξ* = *ξ*_*_ determined by the exponential increase with rate 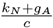 along the unstable fibres of 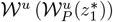, and from *ξ* = *ξ*_*_ onwards determined by the exponential decrease along 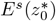, with exponential rates given by the stable eigenvalues 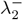 and 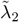.

#### Remark 3

*While the condition r*_*T*_ ≫ 1 *in Theorem 3 is natural (sufficiently strong toxicity feedback induces a JC distribution), the second condition g*_*S*_ *≪*1 *seems less so. Indeed, the necessity for this condition is purely technical, as it allows us to obtain analytical expressions for the stable eigenvalues and eigenvectors of* 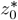 (39). *However, considering the feasible parameter ranges for g*_*S*_, *and in particular for the product g*_*A*_*g*_*N*_ *g*_*S*_, *we infer from Table 2 that g*_*A*_*g*_*N*_ *g*_*S*_ *is small for a significant subset of parameter space – that is, for most values of g*_*A*_ *and g*_*N*_, *the condition that g*_*S*_ *is sufficiently small is not restrictive*.

### 5.5 Wave speed

The analysis of front propagation in excitable media has been a topic of interest for several decades. In his seminal review paper, Van Saarloos [36] used the characterisation *pulled front* for those travelling fronts whose speed is determined by the instability of the spatially homogeneous steady state that is being invaded.

In this section, we analytically determine this ‘linear’ speed a pulled front would have, by a linear analysis near 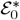. To determine whether the numerically observed fronts can indeed be classified as ‘pulled’, we then compare *c*_*_ with the wave speed computed numerically for the emerging travelling wave solutions.

#### Theorem 4

*The linear wave speed c*_*_ *of a pulled front solution to Equation* (3) *is given by*

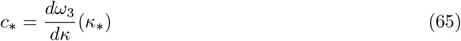

*where ω*_3_(*κ*) ∈ ℂ *is a purely imaginary solution to*

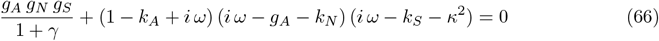

*and* 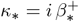 *with* 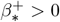 *solution to*

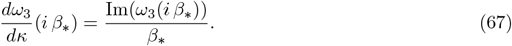

*Proof*. In order to compute the linear wave speed, we analyse the dispersion relation of Fourier modes of the linearisation of Equation (3) at 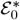. Introducing the diffusion matrix

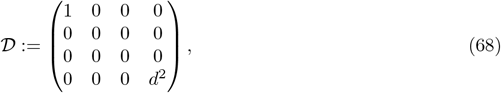

the dispersion relation *ω* = *ω*(*κ*) is given by the solution to

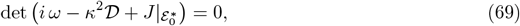

where *ω* ∈ ℂ represents the (generalised) frequency, *κ* ∈ ℂ the (generalised) wave number, and 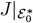 is the Jacobian (21) evaluated at 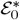. In our case, Equation (69) takes the form of the fourth order polynomial in *ω*

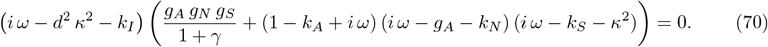

For sake of simplicity, Equation (70) can be equivalently expressed as *ϕ*(*ω, κ*) · *ψ*(*ω, κ*) = 0, where

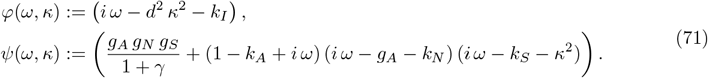

Given a solution *ω*(*κ*) to Equation (70), the linear wave speed *c*_*_ ∈ ℝ and the linear spreading point *κ*_*_ associated to *ω*(*κ*) are given by the solutions to the equations

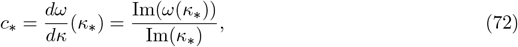

see [18, 36].

Among the four roots *ω*_*i*_(*κ*), *i* = 1, …, 4 satisfying Equation (70) we define *ω*_1_(*κ*) as the unique root of *ϕ*(*ω, κ*) and *ω*_*i*_(*κ*), *i* = 2, 3, 4 as the three roots of the cubic polynomial *ψ*(*ω, κ*). Since we have that *ω*_1_ = −*i* (*d*^2^ *κ*^2^ + *k*_*I*_), we exclude this root from further analysis as *ω*_1_ does not admit solutions to (72).

In our analysis of the other three roots *ω*_*i*_(*κ*), *i* = 2, 3, 4 we introduce the additional assumption (based on our numerical findings, see below) that both *ω* and *κ* are purely imaginary, as the fronts we observe are monotonic, i.e. non-oscillatory, both in space and time, near the trivial steady state 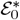. In particular, spatial oscillations around 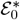 would violate the fundamental model assumption that all model components are non-negative. Hence, we write *κ* = *iβ* with *β* ∈ ℝ. We note that in this case Equation (70) is explicitly solvable, however the analytical expression for *c*_*_ (function of *g*_*A*_ *g*_*N*_ *g*_*S*_, *g*_*A*_ + *k*_*N*_, and *k*_*S*_) is a root of a fifth order polynomial, making it hardly accessible (and hence is not provided here). As observed numerically, two out of three roots *ω*_2_(*κ*) and *ω*_4_(*κ*) are not purely imaginary for every value of *β*, and are therefore further discarded from our investigation. The unique root *ω*_3_(*κ*) is finally used to derive the value of *κ*_*_ = *iβ*_*_ such that Equation (67) holds, i.e.

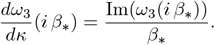

This equation admits two solutions 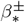 with opposite signs; however, as we are interested in right-moving fronts, we only retain the positive solution 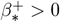. We thus finally obtain the linear wave speed as

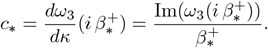

As we do not provide an explicit analytical expression for *ω*_3_(*κ*), in Figure 4 we illustrate a typical plot of the functions 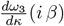 and 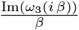 with respect to *β* for the following fixed parameter values (within the ranges reported in Table 2):

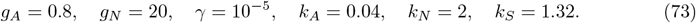

**Figure 4:**
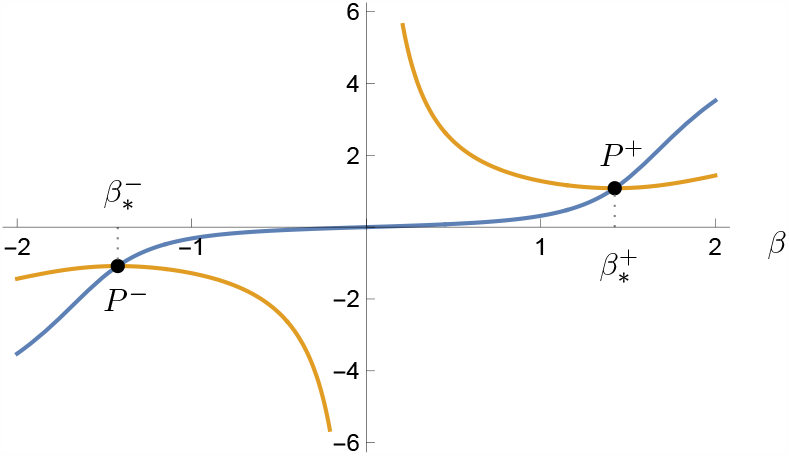
Plot of the functions 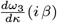 (blue curve) and 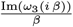 (yellow curve) for parameter values as in Equation (73) and *g*_*S*_ = 0.132. The intersection points between these two curves occurring at 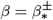 are indicated by the two black points *P* ^*±*^. For these parameter values we observe that 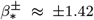 and therefore *P* ^*±*^ = (±1.42, ±1.08).

#### 5.5.1. Numerical investigation

In order to further validate the existence of pulled front solutions travelling with speed *c*_*_ as described in Theorem 4, we perform numerical simulations of System (1) on a one-dimensional domain of size 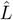 discretized with a spatial grid of 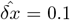 meters including Neumann boundary conditions and the following initial conditions

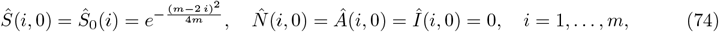

where the dimensional parameters correspond to the ones described in Table 1 and *m* = 3500 corresponds to the number of elements used in the grid. We hence have 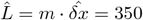 meters. The total simulation time is 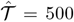 years with timesteps of 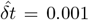 years. Following [33], the numerical scheme used in our simulations is based on a forward Euler integration of the finite-difference equations obtained by discretising the diffusion operator with no-flux (i.e. Neumann) boundary conditions.

The dimensional parameter values fixed in this simulation (other than the ones already fixed in Table 1) are (for unit measures we refer to Table 1)

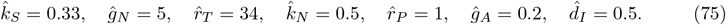

We then investigate two aspects, namely the dependency of the wave speed on the parameter *ĝ*_*S*_ (fixing 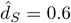) and the dependency of the wave speed on 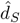 (fixing *ĝ*_*S*_ = 0.033).

A comparison between the values of the linear wave speed obtained from the analytical investigation described in Theorem 4 and the numerical speed computed by means of simulations w.r.t. *g*_*S*_ and *d* is given in Figure 5. To obtain it, we first calculate the dimensional wave speed *ĉ*_*_ as follows, and then derive the nondimensional wave speed as 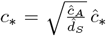. The dimensional numerical wave speed *ĉ* in both scenarios described above – identified by 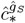 and 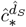, respectively − is obtained by tracking at each time 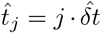 the location of the inflection point in the *Â* profile − defined as 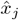 − and subsequently calculating the mean of the difference quotient over a specific range of iterations, namely

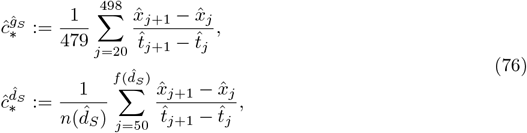

where

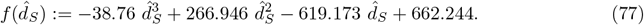

The function 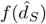 has been derived by interpolating end times in the simulations such that a wave travels with constant shape for 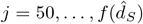. This is due to the fact that the range of 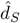 over which the simulation runs has a strong impact on the speed of the travelling wave, which reaches the boundary of the spatial domain sooner for higher values of 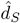. The number of iterations 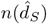 over which the speed 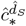 is calculated hence varies with 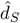. Since, on the other hand, variations of *g*_*S*_ (here intended in its dimensional version) do not exhibit the same properties, the interval over which the numerical wave speed is calculated is here considered as constant.

**Figure 5:**
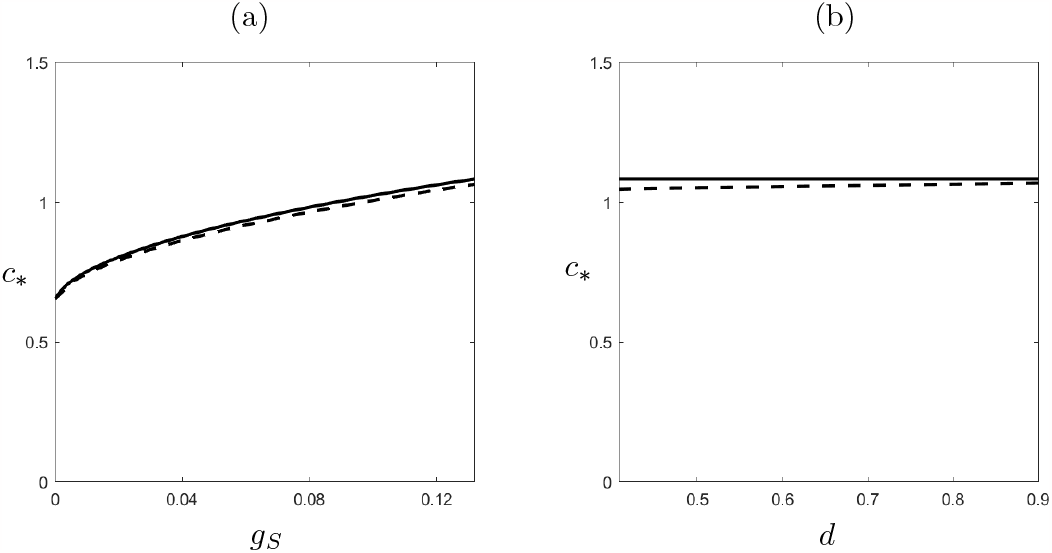
Comparison between the nondimensional wave speed obtained analytically (solid line) and from numerical simulations (dashed line) as a function of (a) *g*_*S*_ with *d* = 0.913 and (b) *d* with *g*_*S*_ = 0.132. The other parameter values are set as in Equation (73) together with *r*_*T*_ = 4080, *r*_*P*_ = 480. We note that these values are obtained by plugging the dimensional values in Equation (75) into Equation (4).

By converting the numerical wave speed in Equation (76) in its nondimensional form 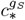 and 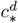, we finally compare it with the analytical values obtained in Theorem 4 (see Figure 5). We note that the strong dependency of the dimensional wave speed 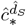 on 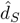 does not imply that the same effect should be valid for the nondimensional speed, which in fact remains approximately constant as *d* varies as shown in Figure 5(b).

We finally observe that the numerical results confirm (up to (10^*−*2^) due to numerical precision) the analytical predictions; such accuracy increases by increasing the size of the domain (by considering larger values of *m*) and thus increasing simulation times as well (see Figure 6). In order to achieve even higher accuracy, the size of the domain should increase with 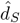 since (as discussed above) for higher values of the seed dispersal coefficient the boundary of the spatial domain is reached sooner by the travelling wave (we note that larger values of 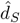 correspond to lower values of its nondimentional counterpart *d*).

**Figure 6:**
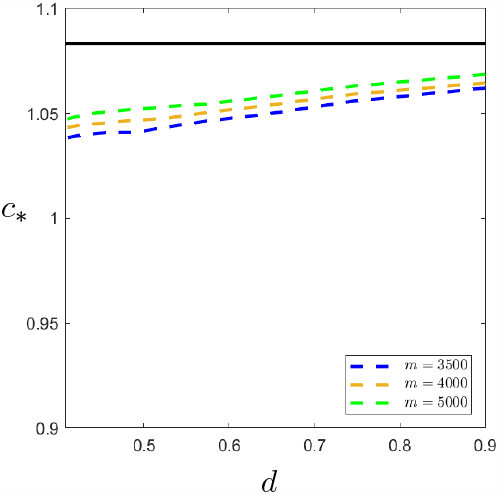
Comparison between the wave speed obtained analytically (solid black line) and from numerical simulations (dashed lined) with the same parameter values as in Figure 5 for different domain sizes 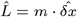 with *m* = 3500 (blue), *m* = 4000 (orange), and *m* = 5000 (green). We note that the accuracy of the numerical wave speed increases with *m*.

## 6. Conclusion

In this work, we have introduced a novel reaction-diffusion-ODE model for (ecologically relevant) transient patterns observed in nature, known as Janzen-Connell distributions. The functional responses adopted in the model, as well as the parameter ranges chosen for the analysis, are based on theoretical assumptions supported by experimental findings. We have included two prominent mechanisms in negative plant-soil feedback - namely growth inhibition and increased mortality - in order to show their key role in the emergence of such transient patterns.

The analytical challenges provided by the complex structure of some functional responses - in particular the germination function - were here overcome by exploiting the small scale of certain parameters in the system. This feature has also played a key role in our thorough investigation of travelling wave solutions, i.e. the theoretical representation of the JC distributions we aimed to describe. Our linear stability analysis allowed us to rule our the existence of Turing(-Hopf) bifurcations and infer the existence of travelling wave solutions for parameter values spanning within ranges of ecological feasibility exhibiting the typical features of JC distributions. Moreover, numerical simulations suggested that the travelling wave solutions admitted by our model in a large area of parameter space correspond to pulled fronts, “pulled” by the linear spreading of small perturbations into the linearly unstable bare soil steady state. The analytical expression for the linear spreading speed was then compared with the numerical speed of one-dimensional waves travelling on a sufficiently large spatial domain – mimicking the unbounded domain of the analytical investigation; the high accuracy revealed by this comparison strongly supports our hypothesis on the pulled nature of the constructed fronts.

As the presented model exhibits a rich and complex structure, several interesting research directions can be further considered. Few examples which we plan to undertake in the future include a deeper investigation of different scenarios corresponding to different combinations of growth inhibition/increased mortality intensity (represented by high/low values of *r*_*T*_ and *r*_*P*_, respectively). Moreover, in order to increase the impact of our model beyond the theoretical sphere, we aim to focus on more realistic ecological scenarios where different trees interact in a limited space (i.e. a bounded domain) and, as a further step, extend our model to a multi-species framework.

## Appendix A Linear stability of 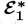 with respect to spatially homogeneous perturbations for *η, ζ* ≠ 𝒪(1)

Based on the values reported in Table 2, and observing that the maximum of 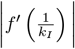 is realised at *I* = *I*_*s*_ and 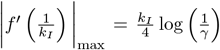, we have that the parameters *η* and *ζ* in (25) can vary within the following ranges:

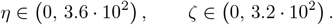

The linear stability of the steady state 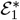 w.r.t. homogeneous perturbations in the case of *η, ζ* ∈ 𝒪(1) has been discussed in the proof of Proposition 2. Here, we look at the other possible regimes, i.e.

I. *η* » 1, *ζ* ∈ 𝒪(1),
II. *η* ∈ 𝒪(1), *ζ* » 1,
III. *η, ζ* » 1.

### A.1. Regime I

*η* » 1, *ζ* ∈ 𝒪(1)

In this case, the dominant term in the characteristic polynomial (26) becomes

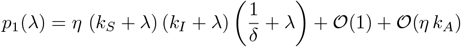

when *λ* ∈ 𝒪(1), which implies that the eigenvalues −*k*_*s*_, −*k*_*I*_, and 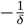 perturb regularly. On the other hand, when |*λ*| » 1 dominant balance gives

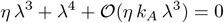

i.e. *λ* = −*η* at leading order. In conclusion, all eigenvalues perturb regularly and remain negative.

### A.2. Regime II

*η* ∈ 𝒪(1), *ζ* » 1

Here, the characteristic polynomial is linear in *ζ* and is given by

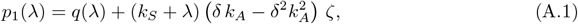

where *q*(*λ*) is a polynomial of degree four in *λ*. By writing 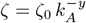, we have that:

- if 0 ≤ *y <* 1, the eigenvalues −*k*_*s*_, −*k*_*I*_, 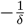, and − (*k*_*N*_ + *η*) perturb regularly;
- if *y* = 1, a regular expansion in *k*_*A*_ leads at leading order to the equation

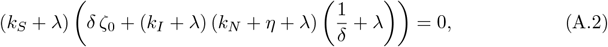

which implies that the eigenvalue −*k*_*S*_ perturbs regularly, while the others shift above by 𝒪(1). These three eigenvalues are negative as long as *ζ*_0_ remains below the Hopf bifurcation value 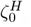, which is found by imposing that the third order polynomial in Equation (A.2) admits a purely imaginary root:

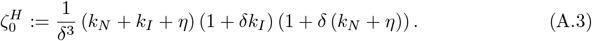

This implies that, for any 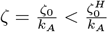, we have three negative eigenvalues.
- if *y >* 1, we have that − *k*_*S*_ is the only eigenvalue which perturbs regularly (i.e. *λ* ∈ 𝒪(1)). On the other hand, dominant balance for |*λ*| » 1 leads to *λ*^3^ = −*δ ζ k*_*A*_ » 1, which implies that here we have one real, negative eigenvalue and two complex conjugates eigenvalues with positive real part for any *ζ* » 1.

In conclusion, in this case we have that all four roots of the polynomial in Equation (A.1) are negative as long as 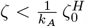.

### A.3. Regime III

*η, ζ* » 1

The characteristic polynomial in this last regime is given by

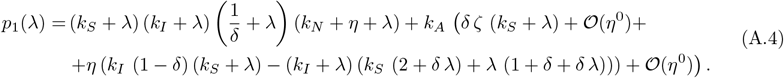

As before, the eigenvalues *λ* = − *k*_*S*_ *<* 0 perturbs regularly, providing one stable, (1) eigenvalue. In order to establish the nature of the other three eigenvalues we need to consider the following scenarios:

- If *η* » *ζ k*_*A*_, the eigenvalues 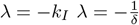 also perturb regularly, providing two negative 𝒪(1) eigenvalues; an additional negative 𝒪(*η*) eigenvalue is given by *λ* = − (*k*_*N*_ + *η*), so in total we have here three stable eigenvalues.
- If *η* ∼ *ζ k*_*A*_, we can write *ζ k*_*A*_ = *ζ*_0_ *η*. Replacing this expression in Equation (A.4) leads to t wo 𝒪(1)eigenvalues with negative real part obtained by solving 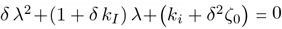 (since (1 + *δ k*_*I*_) *>* 0) and one 𝒪(*η*) eigenvalue *λ* = − *η <* 0. Therefore, in this case we also have three stable eigenvalues.
- If *η* « *ζ k*_*A*_, we have that *δ ζ k*_*A*_ (1 − *δ k*_*A*_) (*k*_*S*_ + *λ*) balances *λ*^4^ +*λ*^3^ (1 − *δ k*_*A*_) *η* + 𝒪(*η*^0^) + 𝒪(*λ*^0^) in Equation (A.4). This leads to further possible scenarios:
  a. If *λ*^4^ » *λ*^3^*η*, the characteristic polynomial at leading order becomes *λ*^4^ + *δ ζ k*_*A*_ *λ* = 0, whose nontrivial solutions consist in two complex roots with positive real part and one negative real root. This implies *λ*^3^ ∼ *ζ k*_*A*_. At the same time, in this case we have *λ* » *η*; these two considerations lead to *ζ k*_*A*_ » *η*^3^.
  b. If *λ*^4^ ∼ *λ*^3^*η*, we can write *λ* = *λ*_0_ *η*; plugging this into the dominant terms of the characteristic polynomial leads to *ζ k*_*A*_ ∼ *η*^3^, hence we can write *ζ k*_*A*_ = *ζ*_0_ *η*^3^. Including this further assumption, the roots of the characteristic polynomial are given by the solutions to 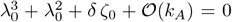. In the case *δ ζ*_0_ = 0, this polynomial admits the negative root *λ* = −1 and a double zero solution. Including the positive term *δ ζ*_0_ hence implies that the negative root perturbs to a root which remains real and negative, whereas the double zero perturbs to a pair of complex conjugate roots with positive real part given by 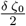 at leading order. In this case no Hopf bifurcation occurs, since the polynomial does not admit purely imaginary roots for any value of *δ ζ*_0_. Therefore, here we have two stable and one unstable eigenvalues.
  c. If *λ*^4^ « *λ*^3^*η*, the characteristic polynomial up to its dominant terms reduces to *λ*^3^ *η* + *δ ζ k_A_ λ* = 0 and is solved by 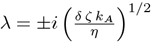, hence requires further unfolding. First, however, we observe that here 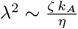, which implies 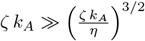 and, in turn, that this scenario corresponds to *ζ k*_*A*_ « *η*^3^. Considering higher order terms leads to the following subcases:
    i. If *η* « *ζ k*_*A*_ « *η*^2^, the characteristic polynomial admits two roots with negative real part given by

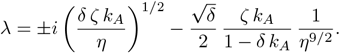
    ii. If *η*^2^ « *ζ k*_*A*_ « *η*^3^, the characteristic polynomial admits two roots with positive real part given by

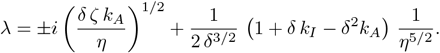
    iii. If *η*^2^ ∼ *ζ k*_*A*_, writing *ζ k*_*A*_ = *ζ*_0_ *η*^2^ leads to the following two roots of the characteristic polynomial with positive real part, given by

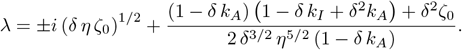

A Hopf bifurcation occurs at

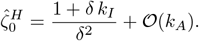
    To summarise, in this case we have that the four eigenvalues of the characteristic polynomial are negative – i.e. 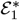 is stable to homogeneous perturbations – as long as 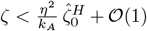. We observe that this value corresponds to the leading order term of 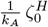 for large *η*, i.e. regime II converges to regime III as *η* → ∞ as expected.

## Acknowledgements

AI is member of Gruppo Nazionale per la Fisica Matematica (GNFM), Istituto Nazionale di Alta Matematica (INdAM). AI acknowledges support from an FWF Hertha Firnberg Research Fellowship (T 1199-N).

## Declaratio of interests

None.

